# Post-mating Refractoriness in *Drosophila melanogaster* Depends Upon Ecdysis Triggering Hormone Signaling

**DOI:** 10.1101/2021.10.29.466485

**Authors:** Matthew R. Meiselman, Anindya Ganguly, Anupama Dahanukar, Michael E. Adams

**Affiliations:** Graduate Program in Cell, Molecular and Developmental Biology, University of California, Riverside, CA, 92521, USA; Department of Molecular, Cell, and Systems Biology, University of California, Riverside, CA, 92521, USA; Department of Entomology, University of California, Riverside, CA, 92521, USA

## Abstract

An individual’s decision to engage in courtship depends on external cues from potential mates and internal cues related to maturation, health, and experience. Hormones allow such information to be conveyed to distal tissues in a coordinated fashion. Here, we show Ecdysis-Triggering Hormone (ETH) is a regulator of male courtship in *Drosophila melanogaster*, and critical for mate choice and courtship inhibition after the completion of copulation. Preventing ETH release increases male-male courtship and decreases post-copulation courtship inhibition (PCCI). Such aberrant male courtship behavior in ETH-deficient males appears to be the consequence of inabilityto integrate pheromone cues into decision making. Silencing of ETH receptor (ETHR) in GR32A-expressing neurons leads to reduced ligand sensitivity and elevated male-male courtship. We find *OR67D* is critical for suppression of courtship after mating, and ETHR silencing in OR67D-expressing neurons, and GR32A-expressing neurons to a lesser degree, elevates post-copulation courtship. Finally, ETHR silencing in the corpus allatum increases post-copulation courtship; treatment of with juvenile hormone analog partially restores normal post-mating behavior. ETH, a stress-sensitive reproductive hormone, appears to coordinate multiple sensory modalities to guide *Drosophila* male courtship behaviors, especially after mating.

## Introduction

Modulation of sensory perception is critical for prioritization of appropriate behaviors under varying physiological conditions. For example, in the fruit fly *Drosophila melanogaster*, mating is costly for both sexes (Chapman et al., 1995; Markow et al., 1978), and the decision to engage in mating behaviors must be weighed with internal state and probability of success (Ejima et al., 2007; Zhang et al., 2019). *Drosophila* males weigh these decisions with a network of *fruitless*-positive neurons (Demir and Dickson, 2005). Reproductive decision-making and consequent behavioral output in male flies is dictated by the activity of P1 neurons (Kimura et al., 2008; Pan et al., 2012). These neurons integrate olfactory, gustatory, visual, and hormonal queues to determine arousal and the ultimate decision to initiate mating (Clowney et al., 2015; Ribeiro et al., 2018; Wu et al., 2019). In addition to direct modulation of the arousal hub, it appears that primary sensory neurons that are necessary for mate perception are also modulated by hormonal state (Lin et al., 2016). This layered control allows the organisms to integrate a variety of factors into their ultimate decision of whether or not to mate.

We recently reported that Ecdysis Triggering Hormone (ETH) persists into the adult stage and acts as a stress hormone in the female, dramatically altering reproductive state (Meiselman et al., 2017; Meiselman et al., 2018). Here, we show that blocking ETH release or elimination of Inka cells, the sole source of ETH, relieves post-copulation courtship inhibition (PCCI) and increases male courtship toward conspecific females and males. Similarly, ETH receptor (ETHR) knock down in aversive pheromone-sensing neurons that express *GR32A* and *OR67D* promotes post-copulation and male-male courtship, suggesting that ETH modulates courtship by affecting pheromone sensitivity. While juvenile hormone (JH) levels are partially responsible for post-copulation courtship inhibition, but treatment with the juvenile hormone analog methoprene (JHA) partially rescues the behavior. We conclude that ETH, a stress-sensitive reproductive hormone, is a potent modulator of courtship, and necessary PCCI.

## Results

### ETH Deficiency Disinhibits Male Courtship Behavior

Normal levels of male courtship behavior depend on JH (Wijesekera et al., 2016), we examined whether ETH-deficient males were impaired in mating efficacy. Surprisingly, we found that ETH deficiency, caused either by expressing the pro-apoptotic protein *reaper* in the Inka cells during the adult-stage (*ETH-Gal4;Tubulin-Gal80*^*ts*^*>UAS-Reaper*)(White et al., 1996), or by adult-specific prevention of ETH release by overexpression of the temperature-sensitive version of the dynamin mutant, *shibire*^*ts*^ (*ETH-Gal4>UAS-shibire*^*ts*^)(Gonzalez-Bellido et al., 2009), causes increased male-female copulation success in a ten-minute window (Figure 1A-B). Injection of ETH peptide to wild-type *Canton-S* males just before being placed with a female dramatically attenuates courtship (Figure 1C). Additionally, both manipulations caused a dramatic increase in courtship toward conspecific males (Figure 1D-E). Most strikingly, we found that ETH-deficient male courtship toward females did not terminate after successful mating, as was observed in control flies (Figure 1G, S1A-B, Supplementary Video 1). In fact, post-copulation courtship in Inka cell-killed males was not significantly different from pre-copulation courtship, though courtship declines dramatically after mating in wild-type, *Canton-S* flies and controls (Figure 1F). This was unlikely to be due to incomplete mating, as copulation duration for ETH-deficient males was significantly longer than controls (Figure S1C-D). We also found that silencing the ecdysone receptor in the Inka cells (*ETH-Gal4;Tub-Gal80ts>UAS-EcR-RNAi*), which reduces ETH expression (Cho et al., 2014), also stimulates post-copulation courtship (Figure S1E).

**Figure 1.**
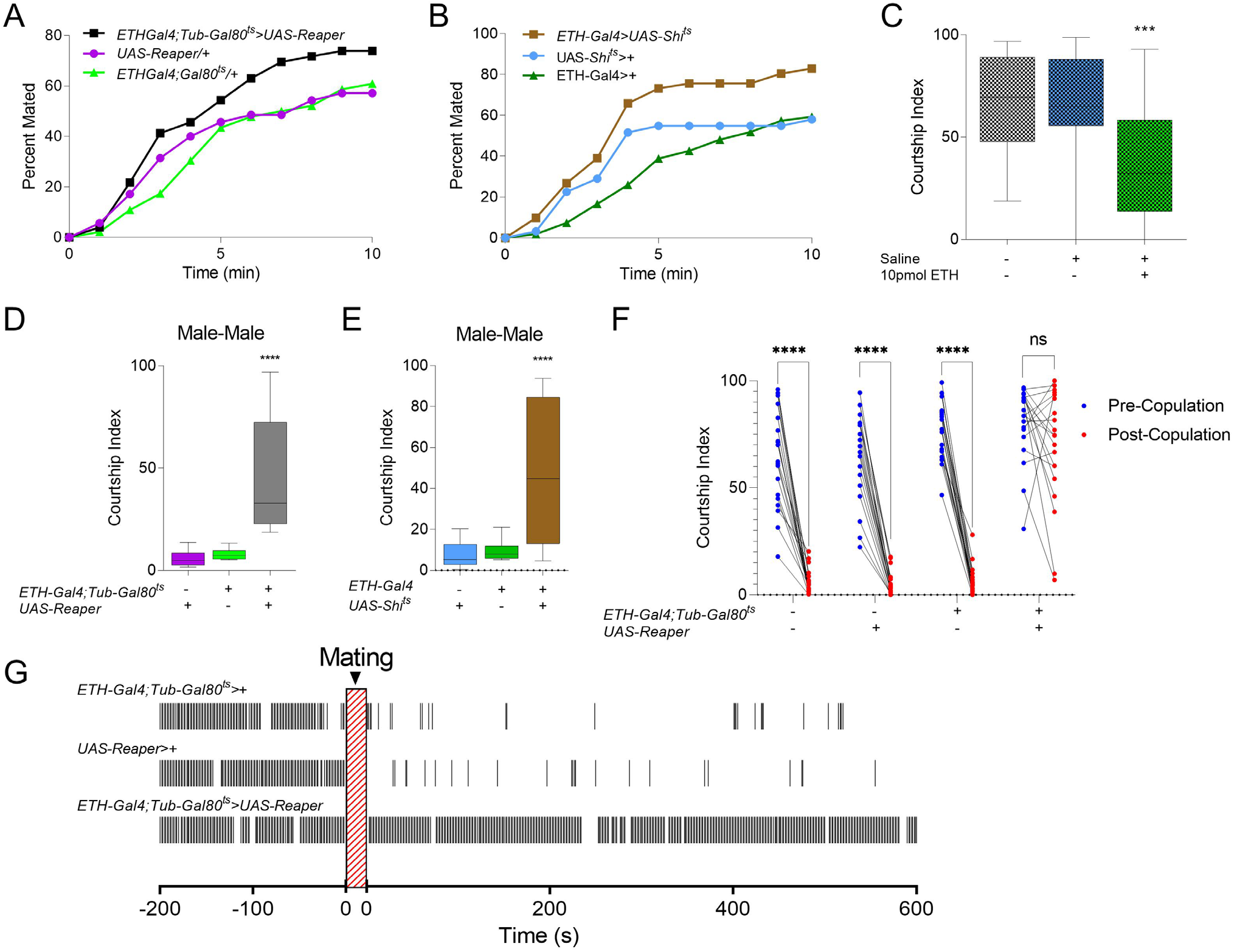
ETH-deficiency increases frequency of courtship behaviors male courtship behavior toward females, males, and mated females. (A-B) Cumulative totals of males successfully mating with virgin *Canton-S* females in ten minutes of indicated genotypes (A) Inka cell-ablated, (*ETH-Gal4;Tubulin-Gal80*^*ts*^*>UAS-Reaper*), (B) Inka cell-Blocked, (*ETH-Gal4>UAS-Shibire*^*ts*^) and genetic controls (n=50-60). (C) Male courtship index toward *wt* females beginning 45 minutes after faux injection (no liquid ejected from capillary), saline injected, or injected with 5 pmol ETH (ANOVA, n=15). (C-D) Male courtship index (time spent courting over total time, 600s) toward *Canton-S* males for Inka cell-ablated (C), Inka cell blocked (D), and genetic controls [One-way analysis of variance (ANOVA), n=15-20]. (E) Time courting 200 seconds before mating and 600 seconds after dismount for an example Inka cell-ablated male and genetic controls. Black bars represent seconds performing courtship behavior. (F) Comparison of courtship index before (Blue dots) and 300 seconds after (Red dots) successful mating for *Canton-S*, Inka cell-ablated males and controls (bootstrap, n=20) ns *p* > .05; ** *p* < .01; *** *p* < .001; **** *p* < .0001.

Male flies typically reduce courtship overtures after mating (Jallon, 1984). This reduction depends upon internal cues from copulation reporting neurons which suppress courtship-promoting NPF neurons (Zhang et al., 2016, 2019), and sensing of external cues which signal female refractoriness (Gillott, 2003; Laturney and Billeter, 2016; Vandermeer et al., 1986). Males deposit pheromones including cis-Vaccenyl Acetate (cVA) and 7-tricosene (7T) on females during copulation, which discourage males from courting during future encounters (Bontonou and Wicker-Thomas, 2014). ETH-deficient males may therefore either be impaired as “senders” (inability to deposit anti-aphrodisiacs) or “receivers” (inability to perceive or integrate information from anti-aphrodisiacs). To clarify, we performed a “Mate-and-Switch” experiment, mating ETH-deficient and wild-type males simultaneously, and swapping their former mates after completion (Figure 2A). We found *Canton-S* (control) males court females formerly mated to ETH-deficient males at low levels, whereas ETH-deficient males courted mated females at high levels (Figure 2B). These findings suggest that ETH modulates internal state to tune courtship. Taken together, ETH levels appear to be critical for maintaining normal courtship behaviors after mating.

**Figure 2.**
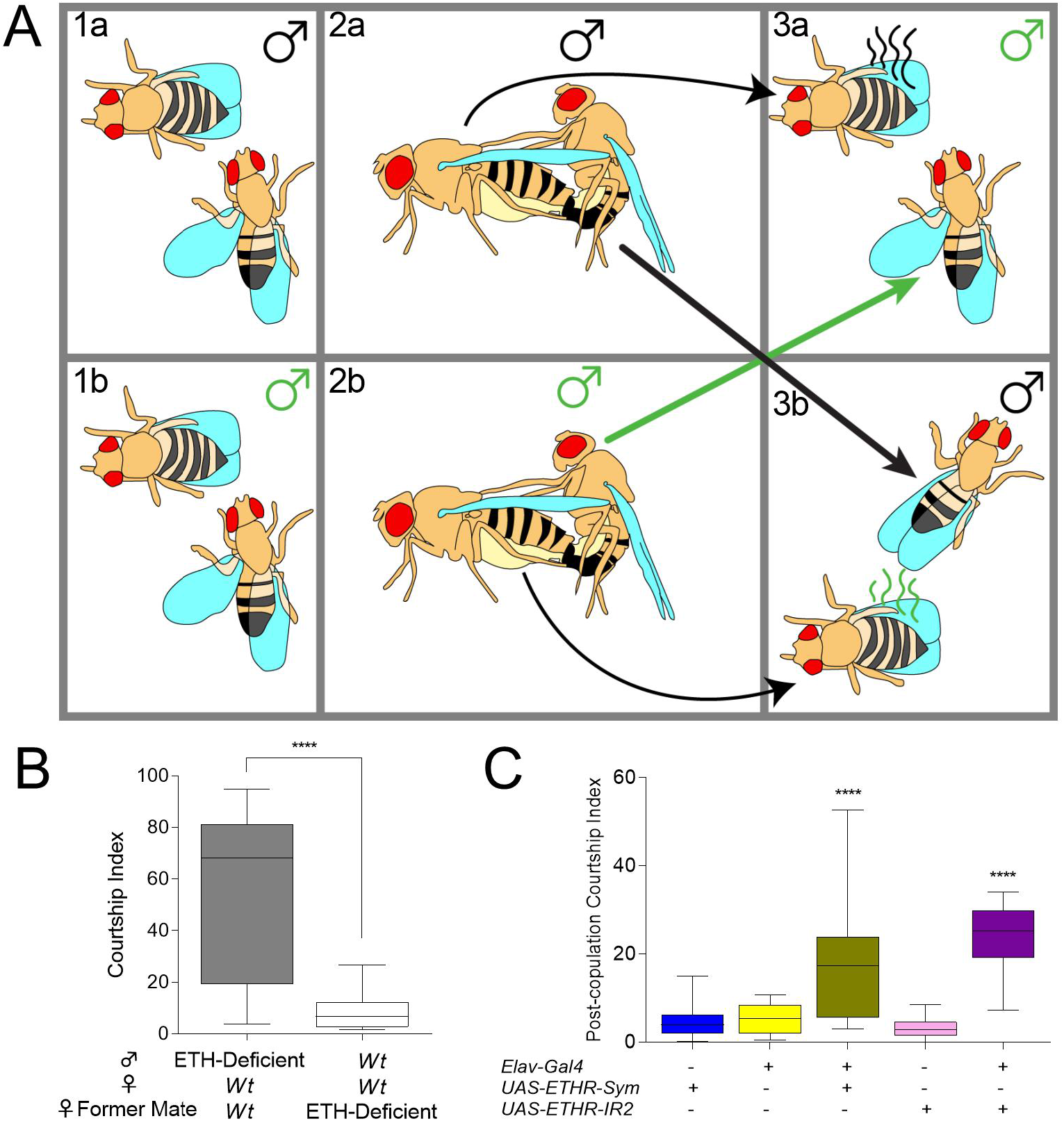
ETH inhibits courtship via perception. (A) Diagram for Mate and Switch experiment. *Wt* and ETH-deficient males court and mate *wt* female counterparts (1a-2a, 1b-2b, respectively). After completion of mating, ETH-deficient males are placed with females mated to *wt* male counterparts (3a), and *wt* males are aspirated and placed with females recently mated to ETH-deficient males from opposite panel (3b). (B) Courtship index after mate and switch for ETH-Deficient and *wt* males after mating and switching (Mann-whitney U, n=15). (C) Male post-copulation courtship index for flies with ETHR knocked down pan-neuronally (*Elav-Gal4>UAS-ETHR-RNAi*) and genetic controls (ANOVA, n=20-21). ns *p* > .05; **** *p* < .0001.

### ETHR-silencing impairs GR32A Neuron Function and Elevates Male-Male Courtship

Males and mated females rely upon pheromones to communicate toward other males that they are an unsuitable mate (Rings and Goodwin, 2019). We recently showed that suppression of male-male courtship depends upon ETH signaling in antennal lobe interneurons and that JH levels and ETH-JH signaling does not impact male-male courtship (Deshpande et al., 2019). ETH-deficient males court conspecific males with more than twice the intensity reported from interneuron-specific ETHR-silencing, we manipulated ETHR expression to determine the extent of ETH modulation of other courtship-related neurons. First, we silenced ETHR pan-neuronally, and observed a significant elevation of post-copulation courtship behavior (Figure 2C). *ETHR-trojan-Gal4* expression is fairly restricted in the brain, labeling Kenyon cells, peptidergic neurons, and olfactory neurons (Lee and Adams, 2021). We identified fruitless-positive, courtship-regulating olfactory neurons innervating VA1lm, VL2A, and DA1 glomeruli, which are postsynaptic targets of OR47B, IR84A, and OR67D, respectively (Figure 3A). *ETHR-Gal4* also labeled taste neurons in the labellum, including those innervating S-type sensilla, which house deterrent neurons that sense bitter compounds and pheromones that inhibit courtship. *ETHR-Gal4* labeled taste neurons in male tarsi but, interestingly, not in female tarsi (Figure 3B-C). Also, *ETHR* and *GR32A* (*ETHR-trojan-Gal4;UAS-mCD8>GR32A-lexA;AOPmCherry*) expression overlaps in the labellum, but not in the tarsi (Figure S2A-B). GR32A, OR47B and OR67D all sense male-derived volatiles (Kurtovic et al., 2007; van Naters and Carlson, 2007; Wang et al., 2011). As ETH-deficient males court other males intensely, we assessed the requirement of ETHR in each of these classes of neurons for inhibition of male-male courtship (Figure 3D). Males with *ETHR* knocked down in GR32A neurons, but not in OR47B or OR67D neurons, significantly increased courtship toward conspecific males, suggesting that ETH may influence GR32A neuron sensitivity.

**Figure 3.**
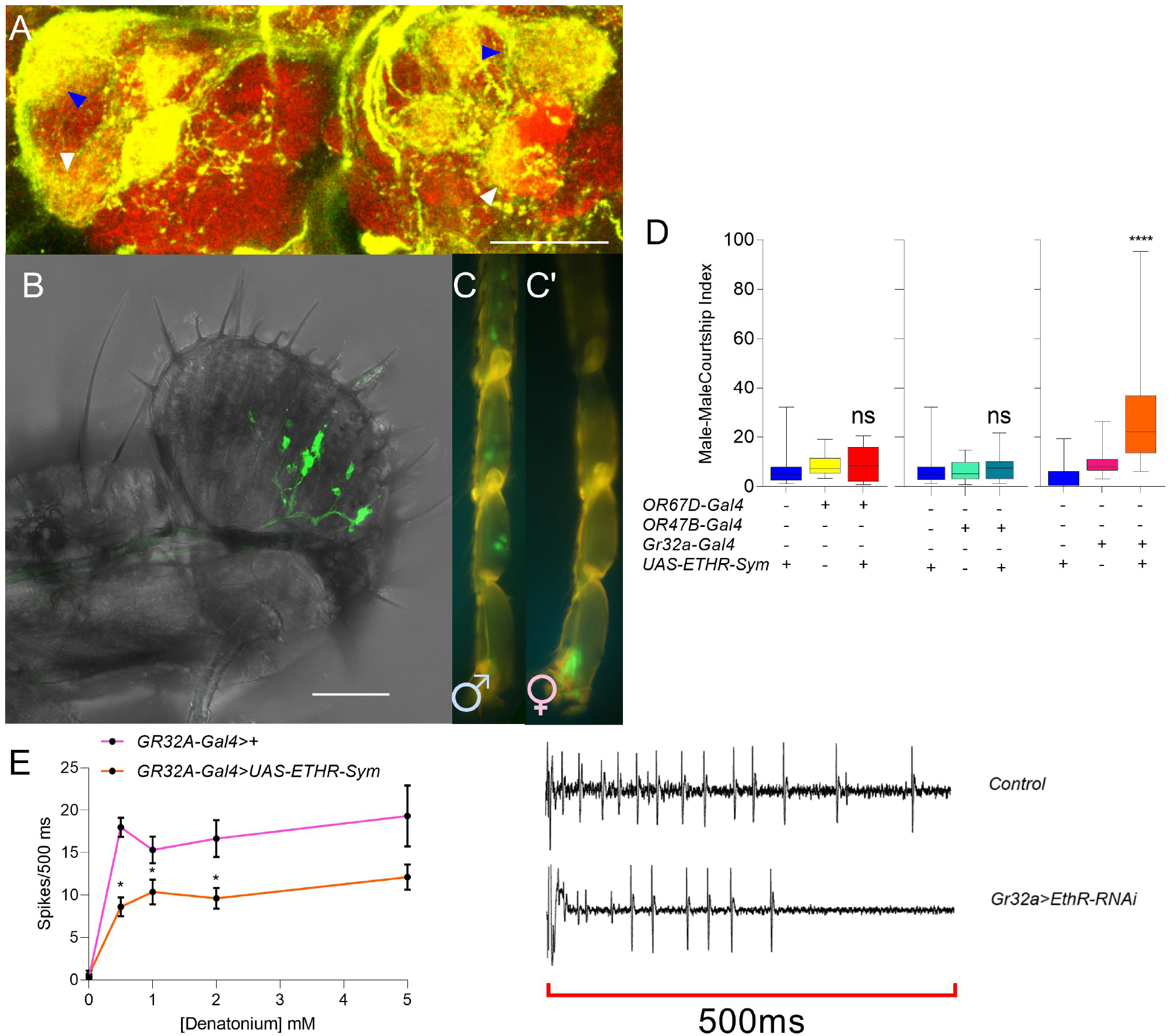
ETHR knockdown reduces sensitivity to ligands in pheromone-sensing neurons. (A) Antennal lobes of *ETHR-trojan-Gal4>UAS-mCD8-GFP* males with arrowheads indicating DA1 glomerulus (targeted by OR67D^+^ neurons, blue) and VA1l/m glomerulus (targeted by OR47B, white)(scale bar 50 μm). (B-C) *ETHR-trojan-Gal4>UAS-mCD8-GFP* expression in male labellum (D), and in prothoracic tarsi (E)(scale bar 50 μm). Sensory hairs are labeled in male tarsi (C), but not females (C’). (D) Male-male courtship index of males with ETHR silenced in pheromone sensory neurons OR67D, OR47B, and GR32A with genetic controls (ANOVA, n=20-25). (E) Extracellular recordings from S6 taste hairs of *GR32A-Gal4>UAS-ETHR-Sym* and *GR32A-Gal4/+* males to 0, 0.5, 1, 2, and 5 mM denatonium for 500 ms after contact (t-test, n=6-8). (F) Example spike for counts. ns *p* > .05; * *p* < .05; **** *p* < .0001.

The overlap in labellar but not tarsal expression of GR32A-LexA and ETHR-Gal4 suggests ETH may target pheromone-sensing neurons in the labellum. GR32A-positive neurons in the labellum are activated by 7T as well as a variety of other bitter tastants (Lacaille et al., 2007; Moon et al., 2009). We tested response to denatonium in ETHR-silenced (*GR32A-Gal4>ETHR-RNAi*) and control males, and found fewer action potentials with a longer latency to response over a range of concentrations (Figure 3E-F). Thus, ETHR deficiency may result in diminished sensitivity to ligands in primary sensory neurons, likely including pheromones.

### ETHR Silencing in OR67D and GR32A-expressing neurons relieves post-copulatory courtship inhibition (PCCI)

Males deposit cVA and 7T on females during mating, which communicate their mated status to future suitors (Laturney and Billeter, 2016). These pheromones are sensed by OR67D and GR32A neurons, whose activity inhibits courtship. We investigated whether these pathways are important for inhibition of courtship immediately after mating. We assessed whether *OR67D*^-^ /- and *GR33A*^-/-^ (a bitter receptor co-expressed with Gr32a and involved in sensing pheromones that inhibit male-male courtship) knock-out mutants were impaired in post-copulation refractoriness (Figure 4A). Only *OR67D*^-/-^ showed elevated courtship after copulation, suggesting that this pathway is indeed a critical component of PCCI. We silenced ETHR in GR32A and OR67D-expressing neurons and, surprisingly, found that ETHR-silencing in both subsets relieves PCCI (Figure 4B). These results suggest that ETH signaling modulates pheromone perception to tune male courtship both before and after mating.

**Figure 4.**
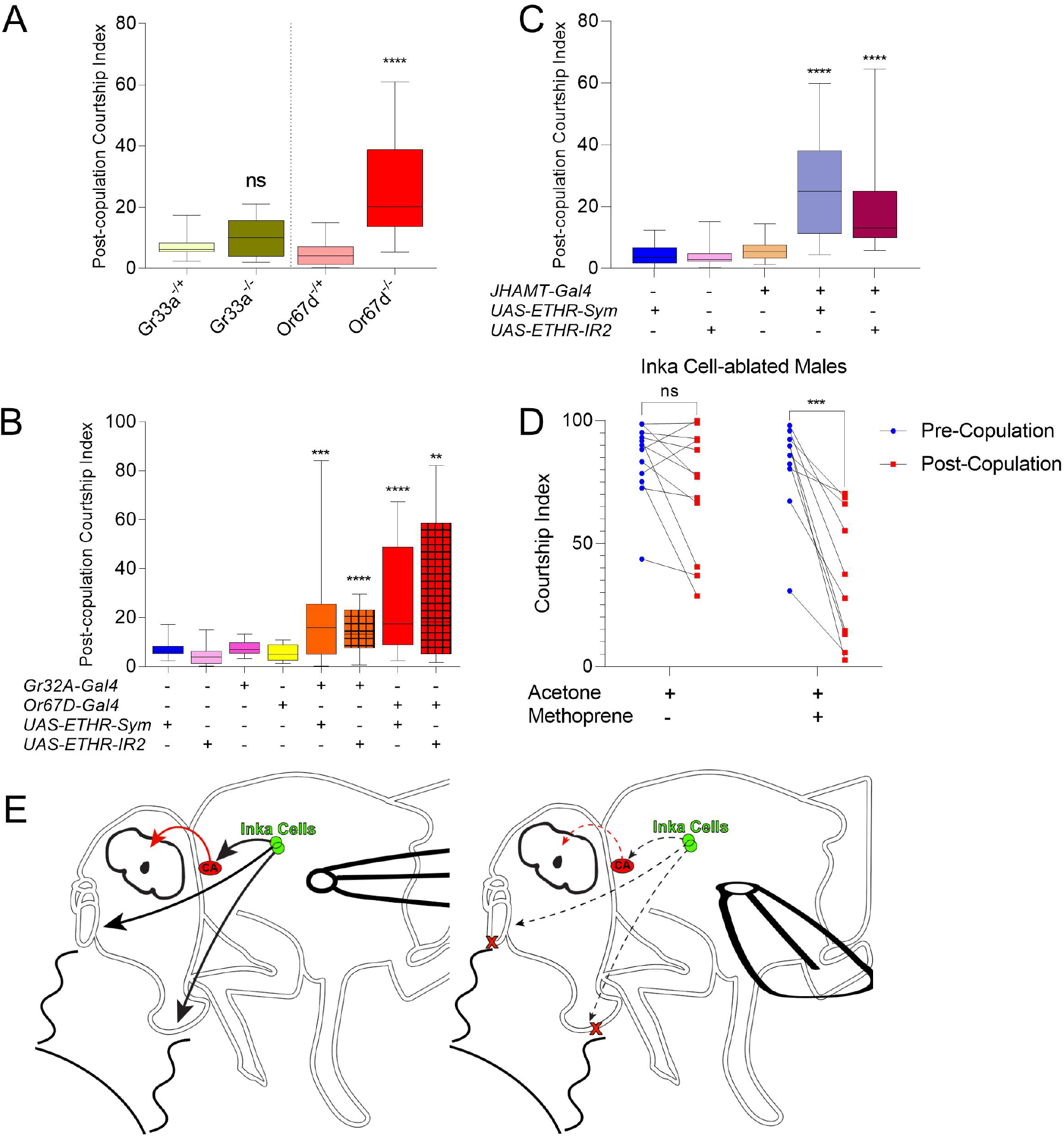
ETH regulates post-copulation courtship by modulation of GR32A^+^ and OR67D^+^ neurons, and JH levels. (A) Post-copulation courtship for GR33A^-/-^ and OR67D^-/-^ and heterozygous controls (Mann-Whitney U, n=10-18). (B) Post-copulation courtship index for *GR32A-Gal4>UAS-ETHR-RNAi, OR67D-Gal4>UAS-ETHR-RNAi* and genetic controls (ANOVA, n=10-25). (C) Post-copulation Courtship for JHAMT-Gal4>UAS-ETHR-RNAi flies and controls. Pre-copulation and post-copulation courtship for Inka cell-ablated (*ETH-Gal4;Tub-Gal80*^*ts*^*>UAS-Reaper*) males treated with either acetone alone or acetone with the Juvenile Hormone Analog, methoprene (n=10-12). (E) Model for the role of ETH in modulation of courtship. In normal conditions (Left), Inka cells (green) release ETH, stimulating Juvenile Hormone synthesis via the Corpus Allatum (Red) and subsequent inhibition of P1 circuitry. Simultaneously, ETH promotes sensitivity to aversive pheromones in peripheral sensory neurons. When ETH is low, the CA and aversive pheromone-sensing neurons are less active, disinhibiting courtship. ns p>0.05; ** *p* < .01; *** *p* < 0.001; **** *p* < .0001.

### Juvenile Hormone Regulates Post-Copulation Courtship

While ligand sensitivity can account for elevated male courtship toward males and mated females, ETH-injected flies reduce courtship toward virgin females significantly, which do not have significant levels of cVA or 7T. ETH-deficient males have low JH levels(Meiselman et al., 2017), and JH inhibits the activity of dopaminergic, pCd, and NPF neurons upstream of P1 courtship command neurons, preventing courtship (Zhang et al., 2021). We therefore sought to determine if JH plays a role in the disinhibition of post-copulation courtship observed in ETH-deficient males. We silenced the ETH receptor (ETHR) specifically in the CA, which reduces total JH levels *(JHAMT-Gal4<UAS-ETHR-RNAi)* (Meiselman et al., 2017). We observed a significant elevation in post-copulation courtship, suggesting JH may also play a role in post-copulation courtship (Figure 4C). We therefore sought to determine whether treatment with the JH analog, methoprene, could restore normal post-mating refractoriness (Figure 4D). Methoprene treatment of ETH-deficient males significantly decreased post-copulation courtship, and restored the normal post-mating decline of courtship, but did not restore courtship to wild-type levels (Figure 4D). JH appears to be critical for post-copulation courtship inhibition, but insufficient to completely account for elevated post-copulation courtship in ETH-deficient males. We therefore propose that ETH inhibits courtship by promoting JH synthesis and increasing sensitivity to courtship-inhibitory pheromones (Figure 4E).

## Discussion

We show here that the peptide hormone ETH is necessary for courtship discrimination and for post-copulation courtship inhibition. We found that ETH-deficient flies show elevated courtship intensity toward same and opposite sex conspecifics, and, strikingly, ETH-deficient males do not terminate courtship after successful copulation. Furthermore, we found that ETH-deficient males are impaired in pheromone sensing or processing, and injection of ETH attenuated courtship intensity toward females. ETHR in *GR32A*-expressing neurons is necessary for normal responses to ligands and prevents abnormal courtship toward conspecific males. Post-copulation courtship is inhibited by *OR67D*, and ETHR-silencing in *OR67D* and *GR32A*-expressing neurons causes disinhibition of post-copulation courtship. Silencing ETHR in the CA also stimulates post-copulation courtship, and treatment of JHA to ETH-deficient males restores inhibition of courtship after mating.

Our data support a model that normal courtship depends upon ETH levels in the hemolymph. At low levels of ETH or with specific inactivation of ETHR, males are less responsive to courtship inhibitory pheromones, both at the level of the primary sensory neuron response and in behavior. JH inhibits courtship in early adulthood (Zhang et al., 2021), and ETH levels are tightly linked with JH. It is therefore likely that ETH targets multiple, semi-redundant signaling systems (GR32A, OR67D, CA) to rapidly or dramatically change courtship levels in response to stimuli. Indeed, ETH is responsible for regulation of reproduction in response to stress, and downstream of ecdysone, a gonadotropic hormone that changes in titer in response to courtship, and can either stimulate or inhibit courtship depending on timing (Ishimoto et al., 2009).

### ETH Signaling in Primary Sensory Neurons

ETH modulates a variety of courtship targets and appears to have a major influence over the drive to mate. Our previous report showed ETHR expression in antennal lobe interneurons is important for mate discrimination (Deshpande et al., 2019). Elimination of ETHR in antennal lobe interneurons caused disinhibition of male-male courtship, presumably through cVA desensitization (Liang et al., 2013; Liu et al., 2011). We show here that knockdown of ETHR in OR67D neurons regulates post-copulation courtship, which is also likely to be cVA-dependent. ETHR silencing in each disinhibits normal courtship, suggesting specific role for ETH in males.

ETHR is a Gq-coupled GPCR, and stimulates calcium release from intracellular stores upon activation (Kim et al., 2006). During the ecdysis sequence, peptidergic ensembles of ETHR neurons mobilize calcium and release peptide signaling molecules that activate centrally patterned behaviors. In this study, excitability of ETHR-expressing chemosensory neurons may be under modulatory influences of circulating ETH, becoming more or less sensitive to pheromone ligands as hormone levels fluctuate, as we observed in GR32A labellar neurons. It is currently unclear over what time frame such hormonal modulation in this context may occur, though during ecdysis, the correspondence between ETH release and behavioral outcomes is a matter of minutes (Kim et al., 2006; Park et al., 2003).

### ETH-JH Signaling Supports Post-Copulation Courtship Inhibition

We recently demonstrated that ETH or ETHR deficiency decreases whole body juvenile hormone levels in both males and females (Meiselman et al 2017). In this work, we show that elimination of ETH or ETHR in the CA elevates courtship after mating. JH has been linked to male reproductive maturation and JH receptors in OR47B neurons promote their receptivity to courtship stimulatory pheromones (Lin et al., 2016). A recent report showed treating mature males with JH suppresses courtship and inhibits calcium activity in courtship-promoting NPF, doublesex-positive pCd neurons, and dopaminergic neurons (Zhang et al., 2021). JH levels do not influence male-male courtship (Deshpande et al., 2019), and post-copulation courtship does not seem ethologically beneficial at any life stage, arguing that this role for juvenile hormone is independent of homeostasis or development. JH targets Kenyon cells to mature and enhance learning memory (Lee et al., 2017; Leinwand and Scott, 2021), and drastically reduced JH levels could prevent flies from forming or accessing a “memory” of a recent, successful mating. Paired with sensory deficit, this may account for total elimination of post-copulation refractoriness.

### Conclusion

Here, we established a relationship between ETH and courtship, wherein ETH stimulates activity in a variety of courtship-inhibiting targets (GR32A-expressing neurons, OR67D-expressing neurons and the CA). We did not investigate how ETH levels may change innately and, in turn, manipulate male courtship. A drop in ETH levels in response to stress would by extension elevate courtship toward conspecifics, though in many other species stress has an inhibitory effect on male libido (Chenoweth, 1981; Lenzi et al., 2003; Marai et al., 2002).

We find the influence of ETH and JH in post-copulation courtship particularly intriguing, as the post-mating change in behavioral state has received less attention than pre-mating courtship. Further investigation is needed to determine how the CRN circuit, which attenuates activity of P1 neurons to induce mating satiety, integrates hormonal changes. After mating, females receive sex peptide, which is released slowly from sperm tails and causes infradian elevation of JH (Liu and Kubli, 2003; Moshitzky et al., 1996; Sugime et al., 2017), allowing females to accelerate oogenesis and replenish egg stores. The male must also replace accessory gland proteins after mating, which depend upon JH for their synthesis (Yamamoto et al., 1988). As males lack circulating sex peptide, this introduces the intriguing possibility that flies employ ETH, an innate allatotropin, to raise JH levels after copulation and inhibit post-copulation courtship. Indeed, in the closely related male carribean fruit fly *Anastrepha suspensa*, JH levels increase after mating (Teal et al., 2000).

We demonstrate here that courtship behavior is critically dependent on ETH signaling, and suggest multiple targets for this dependency. This investigation permits further investigation on how the now well-established neural circuitry of male courtship integrates endocrine state, particularly after mating.

## Acknowledgements

We thank Roscoe Huo, Michael Chitgian, Lola Alade, Michelle Gyulnazaryan and Raymond Tan-Tran for scoring courtship videos, Yuta Mabuchi for comments on the manuscript, Benjamin White and Naoki Yamanaka for fly strains; Anandasankar Ray for fly lines and helpful advice; and the Bloomington Stock Center (NIH P40OD018537) for reagents.

## Author Contributions

M.M. and M.A. conceived the project and designed all the experiments. M.M. carried out all the experiments with undergraduate students mentioned in acknowledgements, with the exception of the electrophysiological recordings, which were designed, carried out, and analyzed by A.G. and A.D. M.M. and M.A. wrote the paper with feedback from all authors.

## Materials and Methods

### Fly Strains

Flies used for immunohistochemistry were raised at 23°C on standard cornmeal-agar media under a 12:12 hr light:dark regimen. Inka cell-ablated flies were raised at the Gal80^ts^ permissive temperature (18°C). Following eclosion, they were moved to the nonpermissive temperature (29°C) for 24 hours, then moved to 23°C until day 4. Inka cell-blocked flies were raised at 18ºC until eclosion, and transferred to 28ºC until day 4. Use of double-stranded RNA constructs for silencing of ETHR (UAS-ETHR-Sym; UAS-ETHR-IR2 line (VDRC transformant ID dna697) were described recently (D.-H. Kim et al., 2015). ETHR-Gal4 was obtained from Ben White (National Institute of Mental Health, Silver Spring). GR32A-LexA, *GR33A*^-/-^, and AOP-mCherry were obtained from Anupama Dahanukar (University of California-Riverside). *OR67D*^-/-^ was obtained from Anandasankar Ray (University of California-Riverside). All other fly lines were obtained from the Bloomington Stock Center (Indiana University, Bloomington, IN): UAS-mCD8-GFP (BS no. 5137), UAS-Reaper (BS no. 8524), Tubulin-Gal80^ts^ (BS no. 7017), ETH-Gal4 (BS no. 51982), OR67D-Gal4 (BS no. 9998), GR32A-Gal4 (BS no. 57622). All flies used for behavior experiments were backcrossed for at least five generations into the Canton-S background.

### Courtship Assays

#### Courtship

For all behavior experiments, males were isolated prior to eclosion in culture tubes with food. Naive day 4 adult male flies were put in a 1cm in diameter courtship chamber with same-aged wild type Canton-S male or virgin female subject and observed. In the case of male-male courtship, test males were scored for stereotypical courtship behaviors and index was created as a measure of time performing those behaviors out of 10 minutes. For male female courtship, time of copulation was recorded, and percentage copulated was assessed for each minute in the first ten-minute interval. Those flies that did copulate were allowed to dismount, and then the male was observed and scored for courtship for the ten minutes immediately following the dismount, which was recorded as post-copulation courtship index. For pre-copulation/post-copulation comparisons, the five minutes (or less if males mated earlier) before mating was compared to only the first five minutes after males and females broke copulation. Time between mounting and dismounting was recorded as copulation duration. Courtship scoring was performed single blind by undergraduates.

#### Competition Assay

Males were placed in a 1.5 cm mating chamber with either a decapitated male and a virgin female and recorded for 20 minutes. Time spent performing courtship behaviors toward each individual was tabulated and represented both as a sum and a percentage of total time spent courting. Preference index was plotted as (Time courting female-Time courting male)/Total time courting either subject.

### Immunohistochemistry

We crossed *ETHR-Gal4* transgenic flies with *UAS-mCD8-GFP* flies to produce progeny expressing GFP in ETHR-expressing cells for immunohistochemical staining. Day 4 adults were dissected in phosphate buffered saline (PBS) and fixed in 4% paraformaldehyde in PBS overnight at 4°C. After washing with PBS-0.5% Triton X-100 (PBST) five times and blocking in 3% normal goat serum in PBST for 30 minutes at room temperature, samples were incubated with anti-NC82 (Sigma, 1:1000) anti-GFP antiserum (Invitrogen, 1:1000) dilution in PBST for 2 days at 4°C. Tissues were washed with PBST three times, incubated with Alexa Fluor 488-labeled goat anti-rabbit IgG (Invitrogen), Alexa Fluor 635-labeled Goat anti-mouse (Life Technologies), 0.5 mg/ml DAPI, and 2% NGS overnight, and washed 4 times for 10 minutes each in PBST before imaging. Immunofluorescence was recorded using a confocal microscope (Leica model SP5) with FITC filter in the Institute of Integrative Genome Biology core facility at UC Riverside.

### Extracellular tip recordings

Extracellular tip-recordings from labellar taste bristles were performed as previously described (Benton and Dahanukar, 2011). All tastants were dissolved in 30 mM tricholine citrate (Sigma, T0252) which functioned as the electrolyte (Wieczorek and Wolff, 1989). Recordings were obtained from male flies aged 4-5 days in 25° C. Neuronal responses were amplified and digitized using an IDAC-4 data acquisition software. Visualization and manual-quantification of the action potential spikes was done by autospike software (Syntech). Number of action potential spikes generated in the first 500 ms following contact artifact was used to quantify the neuronal responses.

### Injections

Males were injected with saline, 5 pmol ETH, or Faux injected using a previously described procedure (Meiselman et al., 2017). In short, males were cold anesthetized and injected at ZT 16:15 with 0 (faux), 50 nl of saline, or 50nl of 100μM ETH, abdominally, from a Drummond Nanoject II. 45 minutes later, each group was placed with a virgin female and scored for courtship for 10 minutes.

### Methoprene Treatment

Within 24 hours of eclosion, adult males or females were cold anesthetized and treated topically on the dorsal side of the abdomen with 36 nl of either acetone, or 0.01% methoprene dissolved in acetone (∼300 nM). The entire procedure took under 20 minutes, after which flies were returned to their housing.

---

**Figure S1.**
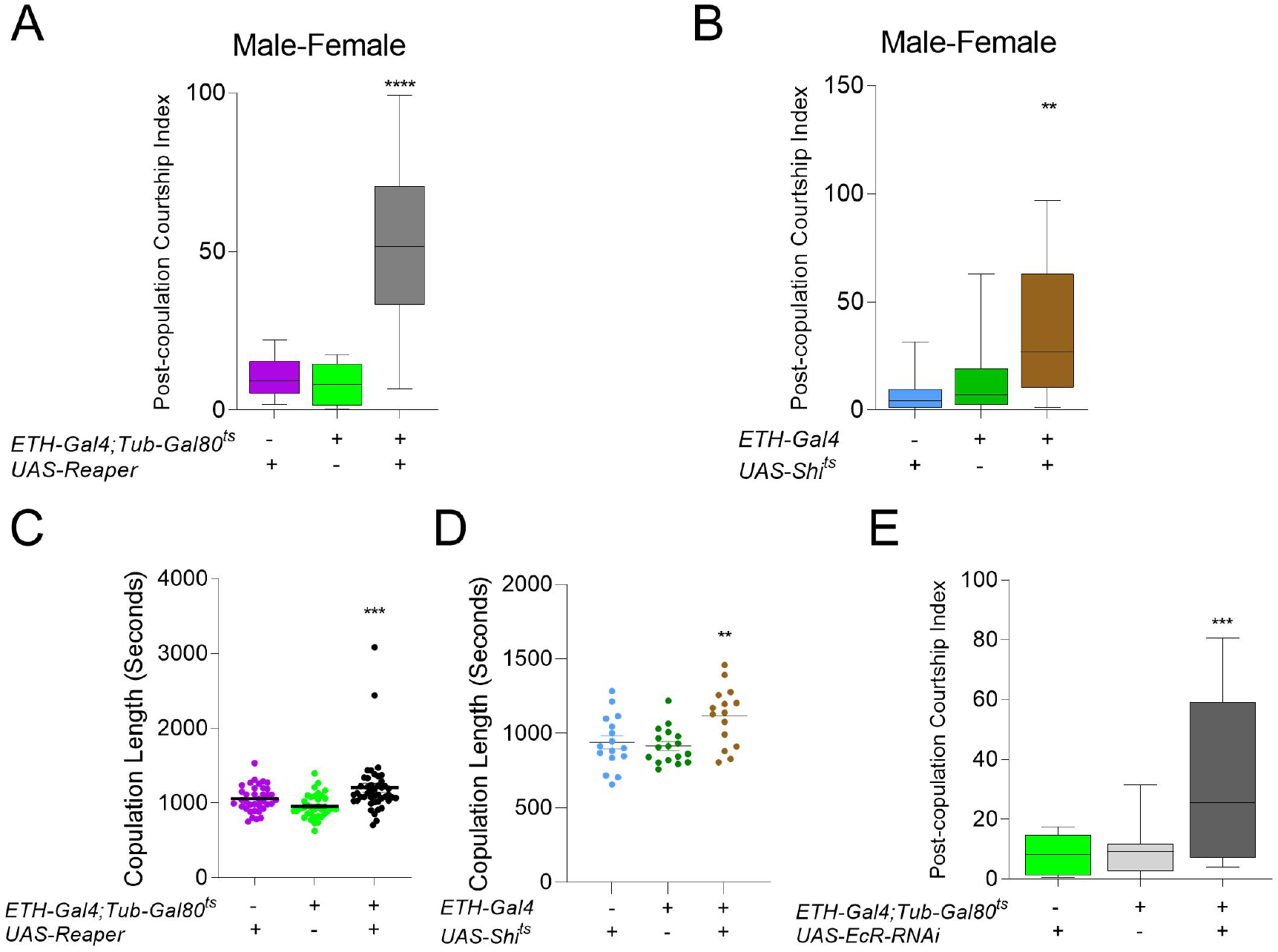
ETH-deficient males exhibit increased copulation length. (A-B) Post-mating courtship index (time spent courting over total time starting at dismounting, 600s) toward a *wt* females for Inka cell-ablated (C), Inka cell blocked (D), and genetic controls (ANOVA, n=20-30). (C-D) Copulation duration (seconds from mounting to dismount) for Inka cell-ablated (*ETH-Gal4;Tub-Gal80*^*ts*^*>UAS-Reaper*)(A, n=30-40) and Inka cell-blocked (*ETH-Gal4>UAS-Shi*^*ts*^)(B, n=15-20). (E) Post-mating courtship index (time spent courting over total time starting at dismounting, 600s) toward a *wt* females with EcR-silenced in Inka cells and genetic controls (ANOVA, n=10-20). ns p>0.05; ** *p* < .01; *** *p* < 0.001; **** *p* < .0001.

**Figure S2.**
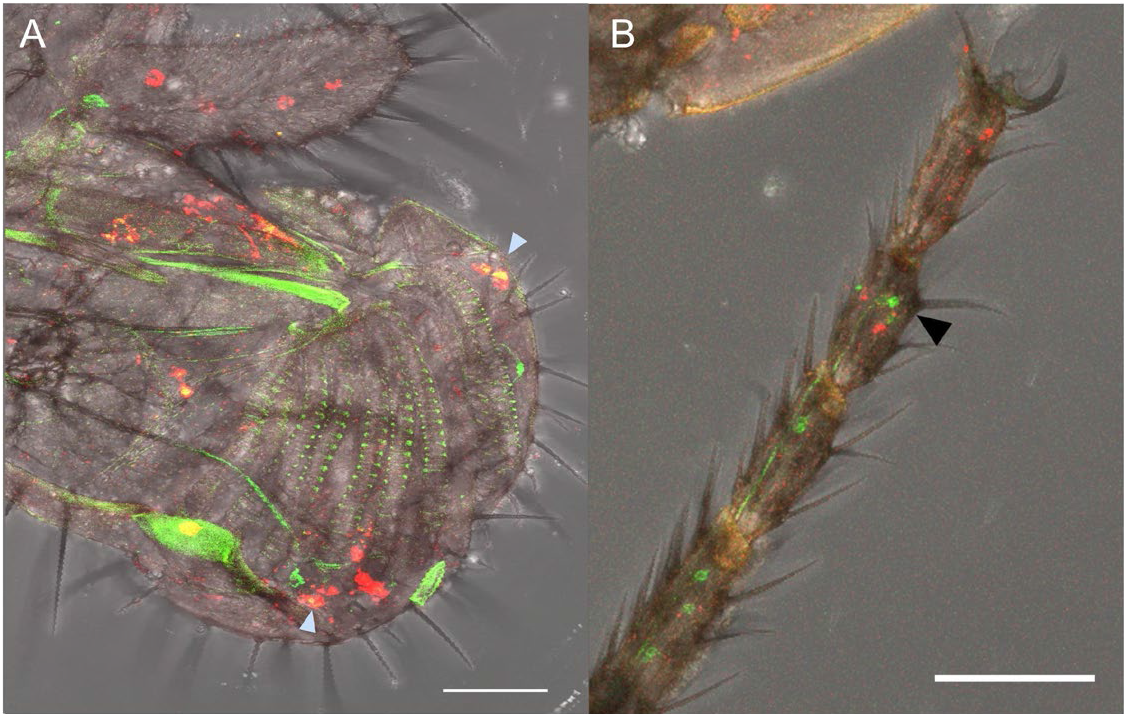
Male expression pattern of ETHR-Gal4. (A-B) *UAS-mCD8;ETHR-Gal4/Aop-mCherry;GR32A-LexA* overlapping expression in labella (A, blue arrowheads) non-overlapping expression in tarsi (B, black arrowhead).

